# Circulating extracellular vesicles in lung cancer patients are not enriched in tumor-derived DNA fragments as revealed by whole genome sequencing

**DOI:** 10.1101/2022.07.22.501161

**Authors:** Norbert Moldovan, Sandra Verkuijlen, Ymke van der Pol, Leontien Bosch, Jan R.T. van Weering, Idris Bahce, D. Michiel Pegtel, Florent Mouliere

**Affiliations:** Amsterdam UMC, Vrije Universiteit Amsterdam, Department of Pathology, Cancer Center Amsterdam, 1081 HV, Amsterdam, The Netherlands; Amsterdam UMC, Vrije Universiteit Amsterdam, Department of Human Genetics and Functional Genomics, Center for Neurogenomics and Cognitive Research, 1081 HV, Amsterdam, The Netherlands; Amsterdam UMC, Vrije Universiteit Amsterdam, Department of Pulmonology, Cancer Center Amsterdam, 1081 HV, Amsterdam, The Netherlands

**Keywords:** Extracellular vesicle, cell-free DNA, whole genome sequencing, cancer

## Abstract

Liquid biopsies contain multiple analytes that can be mined to improve the detection and management of cancer. Beyond cell-free DNA (cfDNA), mutations have been detected in DNA associated with extracellular vesicles (EV-DNA). The genome-wide composition and structure of EV-DNA are poorly characterized, and it remains undecided whether circulating EVs are enriched in tumor signal compared to unfractionated cfDNA.

Here, using whole genome sequencing from selected lung cancer patients with a high cfDNA tumor content (>5%), we determined that the tumor fraction and heterogeneity are comparable between DNA associated with EVs and matched plasma cfDNA. DNA in EV fractions, obtained with standardized size-exclusion chromatography, are comprised of short ∼150-180 bp fragments and long >1000 bp fragments that are poor in tumor signal. Other fractions only exhibit short fragments with similar tumor DNA content. The composition in bases at the end of EV-DNA fragments, as well as their fragmentation patterns are similar to plasma cfDNA. Mitochondrial DNA is relatively enriched in EV fractions.

Our results highlight that cfDNA in plasma is of dual nature, either bound to proteins (including the nucleosome) but also associated to EV. cfDNA associated to small EV (including exosomes) is however not preferentially enriched in tumor signal.

## 1. Introduction

Liquid biopsy provides non-invasive ways to analyze genetic alterations, detect cancer early or monitor residual disease in patients with cancer [1–3]. Cell-free DNA (cfDNA) is the current gold standard for the analysis of genetic alterations with liquid biopsy, but this analyte has clear biological and technical limitations [1, 4]. cfDNA is thought to be released via cell-death mechanisms, the majority through apoptosis [5]. cfDNA might therefore not be released by cancer cells in dormancy, or with slowed proliferating cycle. Moreover, the genetic representation of cfDNA is biased toward shared clones, and might under-represent private mutations [6].

Extracellular vesicles (EV) could complement cfDNA as a liquid biopsy. EVs, and notably endosome derived exosomes can be actively secreted by living cells [7], therefore potentially informing on cell populations underrepresented in cfDNA molecules [8, 9]. Due to this biological difference in their origin, the representation of tumor heterogeneity could differ between EV-DNA and plasma cfDNA [6]. EVs have been reported to contain or be associated to DNA [10, 11].

Prior reports exhibit conflicting results and conclusions on whether EVs contain tumor DNA signals. Approaches based on mutation testing (e.g. digital PCR) have identified either increased or decreased tumor fraction in DNA from EV fraction [12–14]. However, the diversity in the EV isolation and cfDNA analysis methods could explain discordant conclusions. A focus on single loci and mutations in samples with low DNA concentration could be afflicted by high noise to signal ratio and increased stochastic noise in mutation signal. Thus, there is a need to evaluate the characteristics of EV-DNA using sequencing and non-sequencing approaches, and evaluate whether these molecules can be enriched in tumor DNA on a genome-wide scale.

Here we determined the structure and genomic composition of DNA molecules in EV-rich fractions by shallow whole genome sequencing (WGS), and compare them to EV-depleted fractions and unfractionated plasma cfDNA from lung cancer patients. In particular we compared copy number aberrations, size profiles and composition in bases at the end of DNA fragments between the different fractions.

## 2. Materials and Methods

### 2.1. Sample collection

Collection of blood samples in non-small cell lung cancer patients was conducted according to the Declaration of Helsinki and approved by the Amsterdam UMC ethical committee (**Table S1**). Blood was collected in EDTA K2 tubes (BD vacutainer) and plasma isolated within 4 h. Briefly, blood was centrifuged at 900 g for 7 min at room temperature. Supernatant was carefully collected and centrifuged further at 2500 g for 10 min, aliquoted into 0.5 mL tubes and stored at −80°C.

### 2.2. EV isolation

1 mL of plasma per sample was loaded into qEV columns (iZON) with 1.5 mL PBS. This was repeated 3 times, inputting a total of 3 mL plasma per patient. This was done to reach sufficient concentration for sequencing library preparations. Columns were carefully washed between each loading. 20 fractions were further collected using an Izon Automated Fraction Collector (AFC) following the constructor recommendations (**Table S1**). AFC-fractions 1 to 5, 7 to 11, 12 to 15, 16 to 20 were pooled for further analysis.

### 2.3. Electron Microscopy

EV fractions were spotted on carbon/formvar-coated mesh grids. After blotting off the excess liquid, the samples were contrasted by 2% uranylacetate (Polysciences Inc, Cat No 21447-25) in water for 1 minute, the excess stain is blotted off and grids are air-dried. Vesicular structures were imaged with 80 kV Tecnai 12 (ThermoFisher) TEM at 60000x magnification using a 2k x 2k pixel CCD side-mounted camera (Veleta, EMsis GmbH).

### 2.4. Vesicle sizes distribution and concentration

The concentration and diameter of circulating vesicles were determined by tunable-resistive pulse sensing (TRPS), using an Exoid device (iZON). Samples were diluted 50x in an electrolyte buffer and analyzed using a NP250 nanopore at the average result of pressures of 300, 500, and 800 Pa. Concentration and particle size of each pool was determined by measuring calibration beads with a known diameter in nm and concentration in particle/ml and analyzing each sample using the iZON control suite software.

### 2.5. Western blot

Equal volumes of pooled AFC fractions were mixed with sample buffer (for CD63 and CD81 without reducing agent). After boiling for 10 minutes at 95°C, samples were loaded on a 4-15% mini-Protean TGX gel (Bio-RAD). After electrophoresis, proteins were transferred to a 0.2*µ*m nitrocellulose membrane (Amersham) and blocked for 30 minutes with 5% milk in TBS 0.1% Tween at room temperature. Primary antibody incubation was done overnight at 4°C with 5% milk in TBS 0.1% Tween followed by one hour incubation with secondary antibody in 5% milk. Membranes were developed using Pierce ECL western blotting substrate and imaged on a Bio-Rad Gel-Doc XR+ Imager. Primary antibodies used for western blot are anti-CD63 (BD Pharmingen #556019, clone H5C6), anti-CD81 (BD Pharmingen #555675, clone JS-81), anti-Alix (Cell Signaling #2171, clone #3A9), anti-TSG101 (Genetex #GTX70255, clone 4A10), anti-Flotilin 1 (Cell Signaling #18634, clone D32VJ7), anti-Calnexin (Millipore #AB2301). Secondary antibodies used were HRP conjugated rabbit-anti-Mouse IgG (Dako #P260) and HRP conjugated anti-rabbit IgG (Cell Signaling #7074).

### 2.6. DNA isolation

Plasma cfDNA was isolated from all samples using the QIAsymphony Circulating DNA Kit (Qiagen). 3.2 mL of plasma was used for the cfDNA extraction. DNA from EV fractions were isolated using the same protocol. The cfDNA concentration was determined using an Agilent 4200 TapeStation System with the Cell-free DNA ScreenTape Analysis assay (Agilent).

### 2.7. Sequencing preparation

Sequencing libraries were prepared using 1-10 ng of cfDNA and the ThruPLEX® Plasma-seq Kit (Takara Bio) according to the manufacturer’s instructions. Libraries quality was controlled using an Agilent 4200 TapeStation System with the D1000 ScreenTape Analysis assay (Agilent). Libraries were pooled on equimolar amounts and sequenced using 150-bp paired-end runs on the Illumina NovaSeq 6000 using S4 flow-cells (Illumina).

### 2.8. Whole Genome Sequencing analysis

Quality control of demultiplexed data was performed using the FastQC software (v0.11.9). Trimming was done using BBDuk (BBmap v38.79). Quality check post trimming was carried out with FastQC (v0.11.9). Trimmed reads were aligned to human reference genome GRCh38 with BWA-MEM (v0.7.17) using default settings. MarkDuplicates (Picard Tools v2.22.2) was applied to annotate duplicate reads. Reads with a MAPQ score below 5, PCR duplicates, secondary alignments, supplementary alignments and unmapped reads were removed prior to further downstream analysis using samtools (v1.9). Samtools-flagstat software (v1.9) and qualimap software (v2.2.2) were performed as a post alignment quality check.

### 2.9. Copy number analysis

The ichorCNA software (commit 5bfc03e) was used to perform the copy number analysis and estimate the ctDNA tumor fraction [15]. Exceptions to the software’s default settings were as follows: i) An in-house panel of shallow Whole Genome Sequencing (sWGS) normals was created, ii) non-tumor fraction parameter restart values were increased to c(0.95,0.99,0.995,0.999), iii) ichorCNA ploidy parameter restart value was set to 2, iv) no states were used for subclonal copy number and v) the maximum copy number to use was lowered to 3. The reported tumor fraction was retrieved from the data using the highest log likelihood solution.

### 2.10. cfDNA fragment size distribution analysis

The Picard InsertSizeMetrics software (v2.22.2) was used with the default settings on the mapped reads to extract cfDNA fragment size distribution. The proportion of fragments between 20 and 150 bp (P20_150) were calculated from sWGS data for comparison purposes. Plots were constructed in R (v3.6) using the packages ggplot (v3.3.5), dplyr (v1.0.7), tidyr (v1.1.3).

### 2.11. Fragment end sequence composition and diversity

Fragment ends were analyzed using the FrEIA toolkot developed in our group [https://github.com/mouliere-lab/FrEIA.git] controlled by Snakemake (v. 5.14.0). In brief, trimmed and quality-filtered reads were passed to a custom pysam (v. 0.16.0.1) implementation, extracting the first 3 mapped bases from the 5’ end of the reads. Fragments were caracterized based on their first mapping nucleotide and their first three mapping nucleotides. Fractions of these fragment categories were calulated for each sample.

The diversity of the 5’ trinucleotide fragment-end sequences was computed using the Gini index, with the formula:

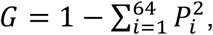

where *P*_*i*_ is the fraction of a given *i* fragment-end sequence.

### 2.12. Statistics

All statistics were calculated in R (v3.6) with the package ggpubr (v0.4.0). P values and statistical test name were included when appropriate in the manuscript and figure captions.

### 2.13. Data availability

Sequencing data will be deposited to the EGA. Other data are available in the supplementary materials.

## 3. Results

### 3.1. DNA in EV-rich fraction is comprised of short and long fragments

To characterize tumor-derived DNA from EV fractions, we selected 6 patients with lung cancer exhibiting a ctDNA fraction >5% and a cfDNA concentration >10 ng/mL plasma based on a prior evaluation [16] (**Figure 1A**). We chose samples with high concentration in cfDNA and relatively high tumor fraction to avoid unfavorable situations where stochastic noise can bias the interpretation of the tumor-derived signal. In addition, 3 mL of plasma was aliquoted for cfDNA analysis using sWGS as previously described (**Figure 1A**).

**Figure 1:**
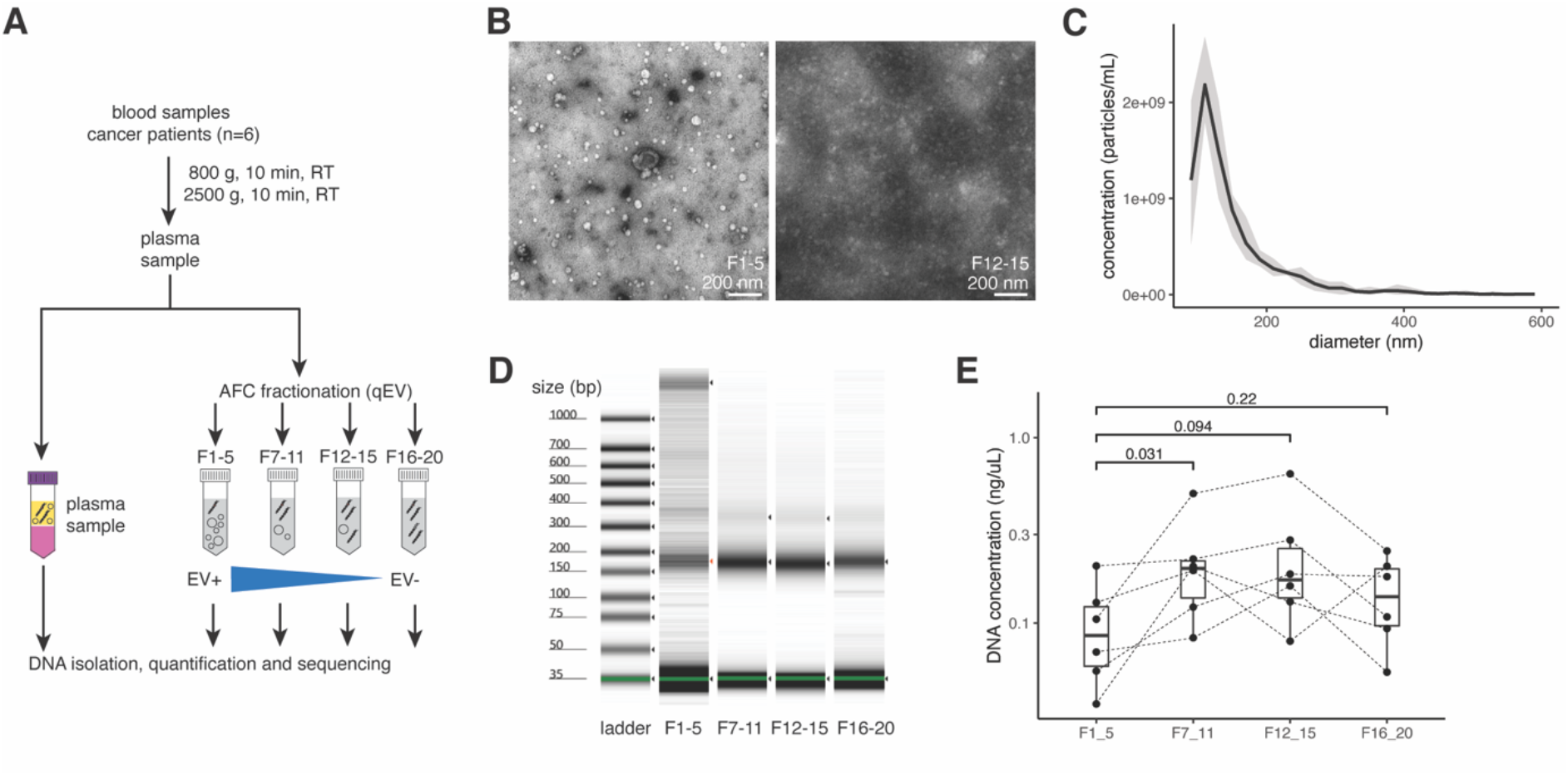
experimental workflow and quality control of the EV separation. **A:** schematic workflow of the AFC fractionation and DNA analysis. **B:** representative EM micrographs of the structures in fraction 1-5 (left panel) and fraction 12-15 (right panel). The white bar indicates the scale (200 nm). **C:** determination of the particle sizes (in nm) and concentration (in particles/mL) using a TRPS device. **D:** concentration (color intensity) and size (bp) of the DNA in the different fractions determined using an electrophoresis device (patient Code53 is shown in this plot). **E:** Concentration in ng/uL of the DNA in the different AFC fractions determined using an electrophoresis device for all patients. P values are indicated (paired Wilcoxon test).

We subjected plasma to size-exclusion chromotography (SEC) with the Automated Fraction Collector (AFC) protocol as indicated by the manufacturer (see Methods). Electron microscopy (EM) revealed vesicle-like membrane structures that range mostly from 50 to 200 nm are enriched in AFC vesicle fractions 1-5 while seemingly absent in protein/HDL fractions 12-15 (**Figure 1B** and **Supplementary Figure 1**). Next, we implemented Western blotting against known markers of small EV (CD63 and CD81) and proteins (see Methods) confirming enrichment in EV in fractions 1-5, and enrichment in proteins in the other fractions (**Supplementary Figure 2**). Particle analysis using Exoid (iZON), based on tunable-resistive pulse sensing (TRPS), confirmed that fractions 1-5 are enriched in small particles with a mean size distribution of ∼144 nm, matching the EM images (**Figure 1C**). Importantly, the concentration in particles of this particular size range is progressively decreased in the other protein-rich fractions (7-11; 12-15; 16-20) (**Supplementary Figure 3**). Together, these data show that EVs can be selectively isolated from plasma samples using the AFC methodology and are enriched in fractions 1-5, and depleted in other fractions.

Automated gel electrophoresis indicates the presence of both short and long DNA in the AFC vesicle fractions 1-5, but only short DNA in the other fractions (**Figure 1D**). Despite this size difference, the observed concentration in DNA is significantly higher in fraction 7-11, compared to the EV-rich fraction 1-5 (p=0.038, paired Wilcoxon test) (**Figure 1E**). The DNA concentration was not significantly different when comparing the fraction 1-5 to the fraction 12-15 and 16-20.

### 3.2. EV-rich fractions are not enriched in tumor DNA

Recovering somatic copy number aberrations could give a potentially more accurate genome-wide determination of the overall tumor fraction of plasma cfDNA samples in comparison to mutation analysis from limited loci [15]. Using sWGS and the ichorCNA software, we determined the tumor fraction in our samples and observed no significant differences between the different AFC fractions (1-5, 7-11, 12-15, 16-20) and the non-fractionated cfDNA plasma sample (paired Wilcoxon test) (**Figure 2A**). Analysis of the difference in the amplitude of the log_2_ratio from the individual genomic bins reveals a correlation between the non-fractionated cfDNA samples, and the different AFC fractions (**Figure 2B**). The different AFC fractions from the same patients cluster together with no apparent difference in the log_2_ratio of the copy number aberrations detected, suggesting that the tumor genomic heterogeneity is represented in a similar fashion in cfDNA, EV-rich and poor fractions (**Figure 2A** and **Supplementary Figure 4**). We also determined that the long DNA (>1000 bp) identified in the EV-rich fractions (1-5) (**Figure 1D**) were exhibiting a significantly lower tumor fraction using sWGS in comparison to the matched cfDNA from plasma samples, or from the other fractions (**Supplementary Figure 5**). Our results indicate that EV-rich fractions (1-5) are not enriched in tumor-derived DNA in comparison to the other AFC fractions or matched total plasma cfDNA.

**Figure 2:**
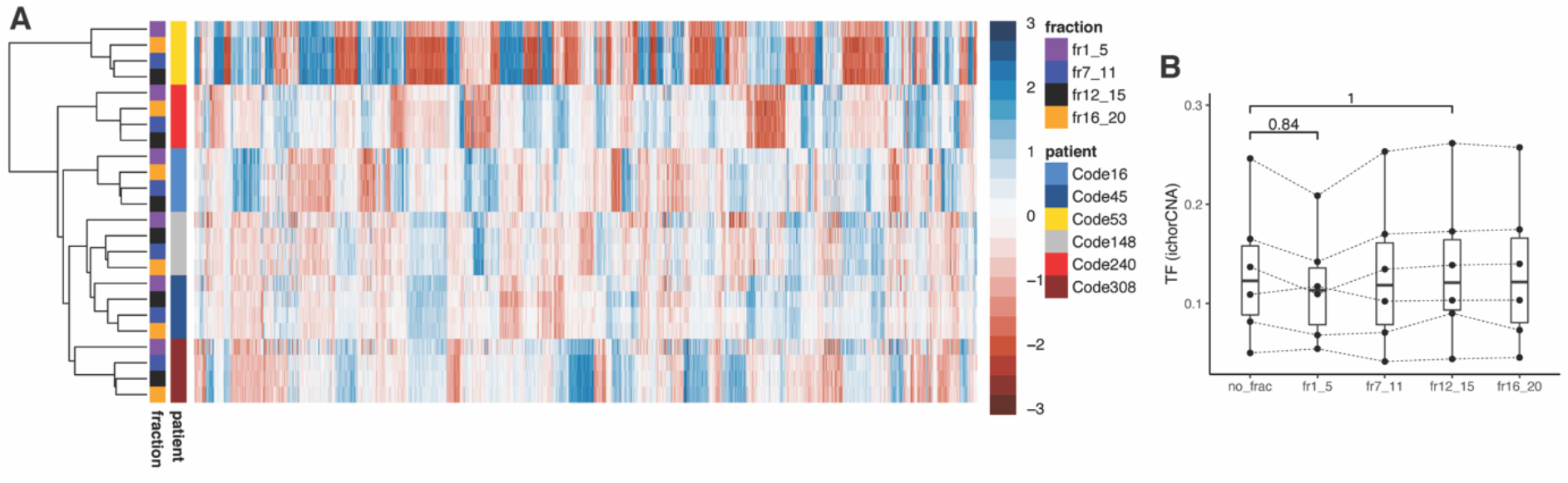
EV-rich fractions are not enriched in tumor-derived DNA. **A:** heatmap of the log_2_ratio copy number aberrations calculated using sWGS data. Rows are samples and columns are genomic positions (100k bins). Blue indicates a relative increase in copy numbers and red a decrease in copy numbers. The respective patients from which the samples are taken, and the type of fractions are indicated as a left annotation. **B:** tumor fraction as estimated from the sWGS data using ichorCNA. P values are indicated (paired Wilcoxon test).

In addition to the copy number aberrations, we analyzed the size of the DNA in the different fractions as it was previously demonstrated it can reflect the presence of tumor-derived cfDNA in plasma [17]. DNA size can be analyzed using the same paired-end sWGS data used for determining their copy number aberrations. We observed in all 4 fractions (1-5, 7-11, 12-15, 16-20) a DNA size distribution with the typical mode at 167 bp previously observed for cfDNA (**Figure 3A** and **Supplementary Figure 6**). The longer DNA fragments are relatively enriched in the EV-rich fractions (1-5) in comparison to the non-fractionated plasma cfDNA and to the other AFC fractions (**Figure 3B** and **Supplementary Figure 7**). The proportion of small DNA fragments between 20 and 150bp (P20_150) are significantly higher in EV-poor fractions (7-11, 12-15, 16-20) than in the EV-rich fractions (1-5), (**Figure 3C**). Beyond the DNA fragmentation, the DNA fragment-end composition was retrieved from the same sequencing data. cfDNA fragment-end in plasma reflects the mechanisms and enzymatic cleavages leading to their release in the circulation [18]. The composition and diversity in trinucleotides at the 5’ end of the DNA fragments reveal no significant differences between the AFC fractions and the non-fractionated plasma cfDNA samples (**Figure 3D** and **Figure 3E**).

**Figure 3:**
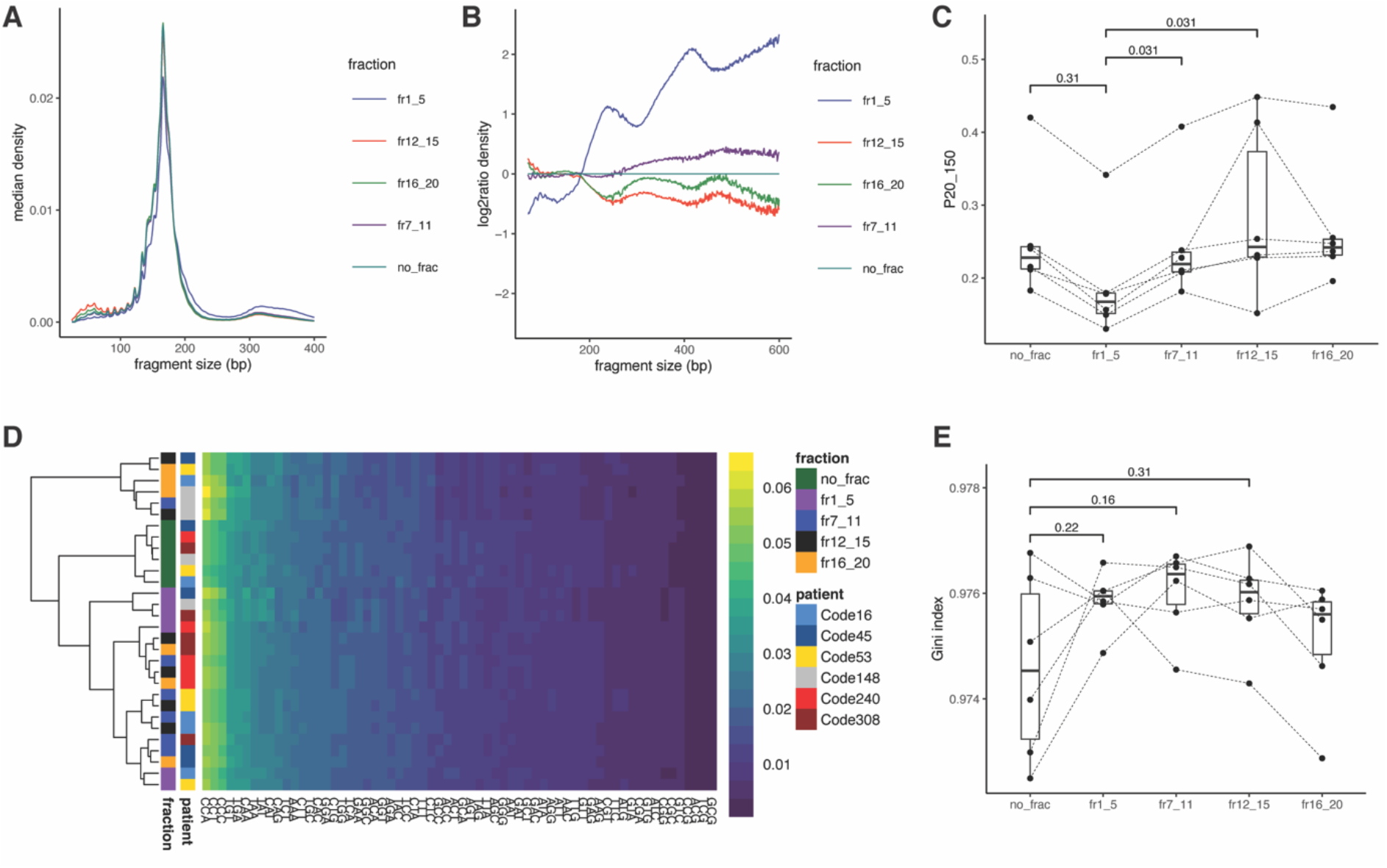
EV-rich fractions are enriched in longer DNA fragments. **A:** median DNA fragment sizes distribution for each of the AFC fractions and matched non-fractionated cfDNA samples. **B:** median log_2_ratio of the DNA fragment size distribution in comparison to the median size distribution of non-fractionated plasma cfDNA samples. **C:** proportion of DNA fragment between 20 to 150bp (P20_150). P values are indicated (paired Wilcoxon test). **D:** heatmap of the proportion of trinucleotides at the end of DNA fragments from the different AFC fractions and matched cfDNA plasma samples. Yellow indicates high proportions and blue low proportions. The respective patients from which the samples are taken, and the type of fractions are indicated as a left annotation. **E:** diversity in the proportion of fragment-end trinucleotides calculated using a Gini index for the DNA of the AFC fractions and matched cfDNA plasma samples. P values are indicated (paired Wilcoxon test).

### 3.3. Non-nuclear DNA are relatively enriched in EV fractions

Beyond nuclear DNA, using untargeted sequencing like sWGS allowed to recover different types of DNA molecules in plasma [19, 20]. We reanalyzed the same dataset by estimating the presence of mitochondrial DNA in the non-fractionated and fractionated samples (**Figure 4**). The fraction of mitochondrial DNA remained a minor component of the total pool of plasma cfDNA (median=0.69 % of the total cfDNA pool). A significant increase in the proportion of mtDNA fragments can be observed in the EV-rich fraction (1-5) (median=3.1%, ∼5 fold relative enrichment), but not in the other fractions (7-11, 12-15, 16-20) (for fr1-5, p=0.031, paired-Wilcoxon test, non-significant for other fractions). Due to their biological proximity with mitochondrial DNA, other non-nuclear DNA in plasma (e.g. bacterial DNA) are likely to be similarly affected in EV-rich fractions [19, 21].

**Figure 4:**
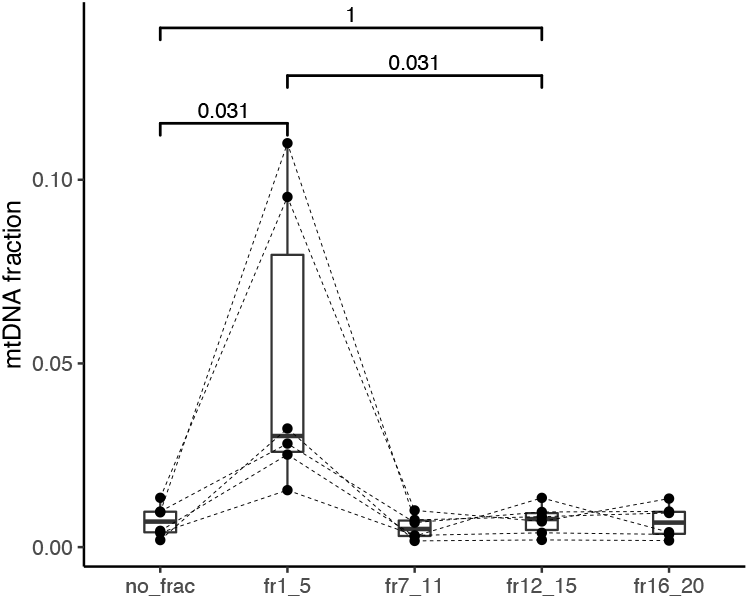
EV-rich fractions are relatively enriched in mitochondrial DNA fragments. The boxplots indicate the proportion of mitochondrial DNA fragments per fractions. P values are indicated (paired Wilcoxon test).

## 4. Discussion

Our results indicate that DNA can be consistently detected in EV-rich fractions (1-5) and other EV-poor fractions (7-11, 12-15, 16-20) collected from the plasma of lung cancer patients using AFC fractionation. The biological properties (fragmentation pattern, size, fragment-end composition) of the DNA identified in the EV-poor fractions are similar to the one observed previously for plasma cfDNA. The EV-rich fraction contains small (∼150-180 bp) and large DNA (>1000 bp), the small DNA exhibiting similar biological properties to plasma cfDNA. This suggests that a fraction of cfDNA in the bloodstream of patients with lung cancer is associated with EV (either inside or bound to EV).

Prior reports indicate that mutations could be more easily detected in exosomes or EV from plasma samples than in cfDNA [12, 13, 22–24]. The observed variation in the ctDNA fractions and detection rates reported in prior reports can be explained by the differences in the EV isolation methods, non-standardized EV characterization post-fractionation, the diversity of methods used for detecting tumor signal, or issues related to the stochastic noise for detecting molecules with mutations in low DNA concentrations. Here, by selecting cancer samples with previously known high tumor fraction and high concentration in DNA, as well as by using state of the art EV fractionation and characterization methods, and genome-wide DNA sequencing method, we by-passed these sources of potential biases. We observed that on a genome-wide scale there is no enrichment in tumor signal in the DNA associated to EV-rich fractions, in comparison to EV-poor fractions or non-fractionated plasma cfDNA. An analysis of the copy number aberrations from the sWGS indicate no significant differences between the copy number aberration profile of cfDNA and EV-rich or EV-poor fractions, suggesting a similar origin for the DNA in these different samples (**Figure 2A**).

Our conclusions are limited by the small cohort of cases included in this study (n = 6). Nevertheless, we only selected lung cancer cases with a high tumor fraction (>5%) clearly detectable using sWGS based on prior results [16], to draw clear conclusions regarding the genome-wide content of the EV-DNA and matched cfDNA samples. We focused our analysis to lung cancer samples, and cannot conclude that the tumor content in EV-DNA might differ in other cancer types. The tumor content greatly differs in cfDNA from various cancer type [25], notably for brain and renal malignancies, and we could anticipate similar differences for EV-DNA. Finally, the genome-wide content of EV-DNA might differ depending on the fractionation method due to the diversity of EV [26]. In particular methods based on size-exclusion chromatography (like AFC) have the advantage to recover purer fractions of EV, but are biased towards smaller EV [27]. Large EV might contain specific DNA populations and will require a different purification approach to formally conclude on their genome-wide tumor content [12]. We did not perform DNAse and/or proteinase treatment of the plasma fractions [28]. We cannot exclude that the EV-DNA in the lumen which is protected from external DNAses underlies the dual length distribution.

In summary, cfDNA in plasma is of dual origin, either bound to proteins (including the nucleosome, as canonically reported) and also associated to EV. cfDNA associated to small EV (including exosomes) is not preferentially enriched in tumor signal.

## 5. Acknowledgements

The authors are thankful to the Amsterdam UMC Liquid Biopsy Center for the logistical support and advices. Y.P., and F.M. are funded by the Amsterdam UMC Liquid Biopsy Center, an initiative made possible through the Stichting Cancer Center Amsterdam. N.M. and F.M. are supported by a Dutch Cancer Fund (KWF-12822). The EM at the VU campus is supported by NWO (91111009). This work was carried out on the Dutch national e-infrastructure with the support of SURF Cooperative.

## 7. Contributions

Conceptualization: F.M. Data curation: F.M. Formal Analysis: F.M.

Funding acquisition: D.M.P. and F.M.

Investigation: N.M., S.V., Y.P., L.B., and J.W.

Methodology: N.M., S.V., Y.P., L.B., and F.M.

Project administration: F.M.

Resources: I.B, D.M.P., and F.M.

Software: N.M., Y.P and F.M.

Supervision: D.M.P, and F.M.

Validation: F.M.

Visualization: F.M.

Writing – original draft: N.M. and F.M.

Writing – review & editing: N.M., S.V., Y.P., L.B., J.W., I.B., D.M.P., and F.M.

## References

1. Heitzer E, Haque IS, Roberts CES, Speicher MR. Current and future perspectives of liquid biopsies in genomics-driven oncology. Nat. Rev. Genet. 2019; 20(2):71–88.

2. van der Pol Y, Mouliere F. Toward the Early Detection of Cancer by Decoding the Epigenetic and Environmental Fingerprints of Cell-Free DNA. Cancer Cell 2019; 36(4):350–368.

3. Wan JCM, Massie C, Garcia-Corbacho J et al. Liquid biopsies come of age: Towards implementation of circulating tumour DNA. Nat. Rev. Cancer 2017; 17(4):223–238.

4. Merker JD, Oxnard GR, Compton C et al. Circulating Tumor DNA Analysis in Patients With Cancer: American Society of Clinical Oncology and College of American Pathologists Joint Review. J. Clin. Oncol. 2018; 36(16):1631–1641.

5. Jahr S, Hentze H, Englisch S et al. DNA fragments in the blood plasma of cancer patients: Quantitations and evidence for their origin from apoptotic and necrotic cells. Cancer Res. 2001; 61(4):1659–1665.

6. Murtaza M, Dawson S-J, Pogrebniak K et al. Multifocal clonal evolution characterized using circulating tumour DNA in a case of metastatic breast cancer. Nat. Commun. 2015; 6(1):8760.

7. Jeppesen DK, Fenix AM, Franklin JL et al. Reassessment of Exosome Composition. Cell 2019; 177(2):428-445.e18.

8. Pegtel DM, Gould SJ. Exosomes. Annu. Rev. Biochem. 2019; 88:487–514.

9. Drees EEE, Roemer MGM, Groenewegen NJ et al. Extracellular vesicle miRNA predict FDG-PET status in patients with classical Hodgkin Lymphoma. J. Extracell. Vesicles 2021; 10(9):e12121.

10. Yokoi A, Villar-Prados A, Oliphint PA et al. Mechanisms of nuclear content loading to exosomes. Sci. Adv. 2019. doi:10.1126/SCIADV.AAX8849/SUPPL_FILE/AAX8849_SM.PDF.

11. Takahashi A, Okada R, Nagao K et al. Exosomes maintain cellular homeostasis by excreting harmful DNA from cells. Nat. Commun. 2017 81 2017; 8(1):1–16.

12. Vagner T, Spinelli C, Minciacchi VR et al. Large extracellular vesicles carry most of the tumour DNA circulating in prostate cancer patient plasma. J. Extracell. Vesicles 2018. doi:10.1080/20013078.2018.1505403.

13. Hagey DW, Kordes M, Görgens A et al. Extracellular vesicles are the primary source of blood-borne tumour-derived mutant KRAS DNA early in pancreatic cancer. J. Extracell. Vesicles 2021; 10(12):e12142.

14. Allenson K, Castillo J, San Lucas FA et al. High prevalence of mutant KRAS in circulating exosome-derived DNA from early-stage pancreatic cancer patients. Ann. Oncol. 2017; 28(4):741–747.

15. Adalsteinsson VA, Ha G, Freeman SS et al. Scalable whole-exome sequencing of cell-free DNA reveals high concordance with metastatic tumors. Nat. Commun. 2017; 8(1):1324.

16. Moldovan N, van der Pol Y, van den Ende T et al. Genome-wide cell-free DNA termini in patients with cancer. medRxiv 2021:2021.09.30.21264176.

17. Mouliere F, Chandrananda D, Piskorz AM et al. Enhanced detection of circulating tumor DNA by fragment size analysis. Sci. Transl. Med. 2018; 10(466):eaat4921.

18. Lo YMD, Han DSC, Jiang P, Chiu RWK. Epigenetics, fragmentomics, and topology of cell-free DNA in liquid biopsies. Science (80-.). 2021. doi:10.1126/SCIENCE.AAW3616.

19. Burnham P, Kim MS, Agbor-Enoh S et al. Single-stranded DNA library preparation uncovers the origin and diversity of ultrashort cell-free DNA in plasma. Sci. Rep. 2016; 6(1):27859.

20. Ma M-JL, Zhang H, Jiang P et al. Topologic Analysis of Plasma Mitochondrial DNA Reveals the Coexistence of Both Linear and Circular Molecules. 2019. doi:10.1373/clinchem.2019.308122.

21. Tulkens J, De Wever O, Hendrix A. Analyzing bacterial extracellular vesicles in human body fluids by orthogonal biophysical separation and biochemical characterization. Nat. Protoc. 2019 151 2019; 15(1):40–67.

22. Yu W, Hurley J, Roberts D et al. Exosome-based liquid biopsies in cancer: opportunities and challenges. Ann. Oncol. 2021; 32(4):466–477.

23. Castellanos-Rizaldos E, Grimm DG, Tadigotla V et al. Exosome-based detection of EGFR T790M in plasma from non–small cell lung cancer patients. Clin. Cancer Res. 2018; 24(12):2944–2950.

24. Allenson K, Castillo J, San Lucas FA et al. High prevalence of mutant KRAS in circulating exosome-derived DNA from early-stage pancreatic cancer patients. Ann. Oncol. 2017; 28(4):741–747.

25. Bettegowda C, Sausen M, Leary RJ et al. Detection of circulating tumor DNA in early- and late-stage human malignancies. Sci. Transl. Med. 2014; 6(224):224ra24.

26. Xu R, Rai A, Chen M et al. Extracellular vesicles in cancer — implications for future improvements in cancer care. Nat. Rev. Clin. Oncol. 2018; 15(10):617–638.

27. Van Deun J, Mestdagh P, Agostinis P et al. EV-TRACK: Transparent reporting and centralizing knowledge in extracellular vesicle research. Nat. Methods 2017; 14(3):228–232.

28. Neuberger EWI, Hillen B, Mayr K et al. Kinetics and Topology of DNA Associated with Circulating Extracellular Vesicles Released during Exercise. Genes 2021, Vol. 12, Page 522 2021; 12(4):522.

