## Supplementary Figures for "Circulating extracellular vesicles in lung cancer patients are not enriched in tumor-derived DNA fragments as revealed by whole genome sequencing"

**Supplementary Figure 1:**

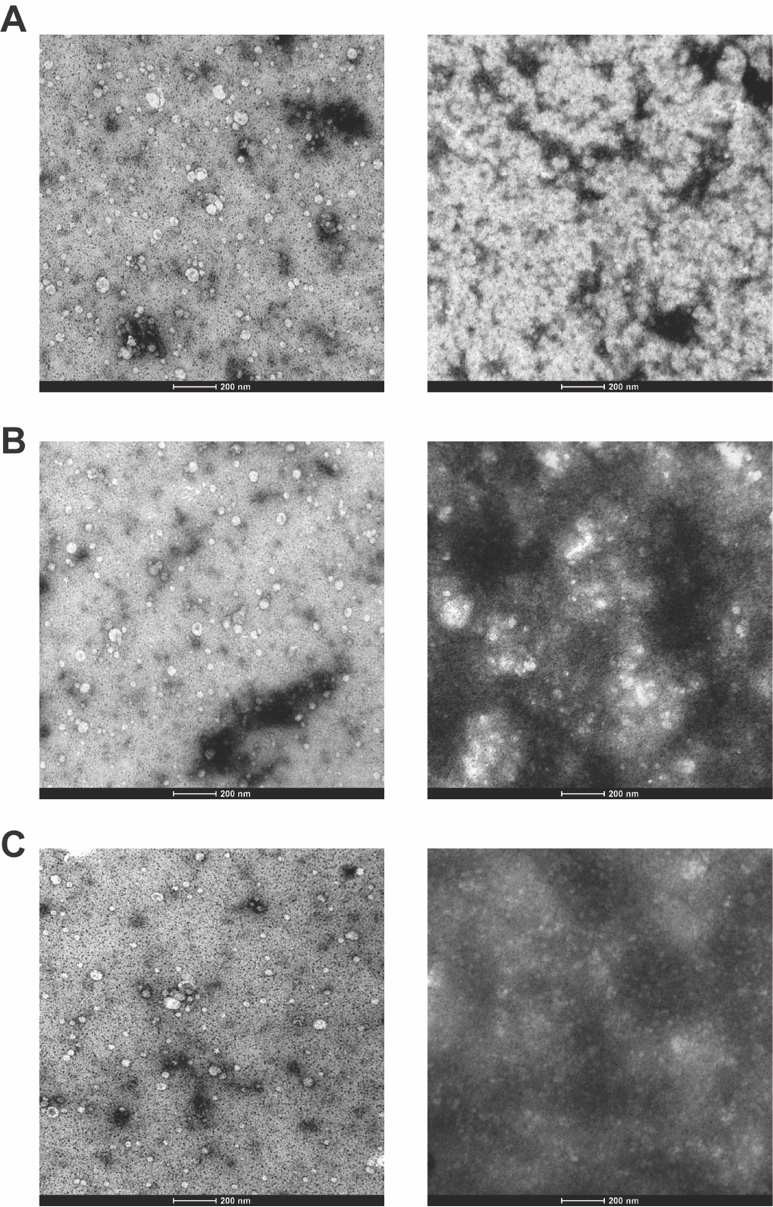

**Supplementary Figure 1:** EM pictures from AFC fractions 1-5 (left panels) and AFC fractions 12-15 (right panels). 10 pictures were taken per sample and one was randomly selected in this plot. Pictures were taken from 3 patients in total. Scale bar indicates 200 nm. **A:** patient Code53. **B:** patient Code148. **C:** patient Code308.

**Supplementary Figure 2:**

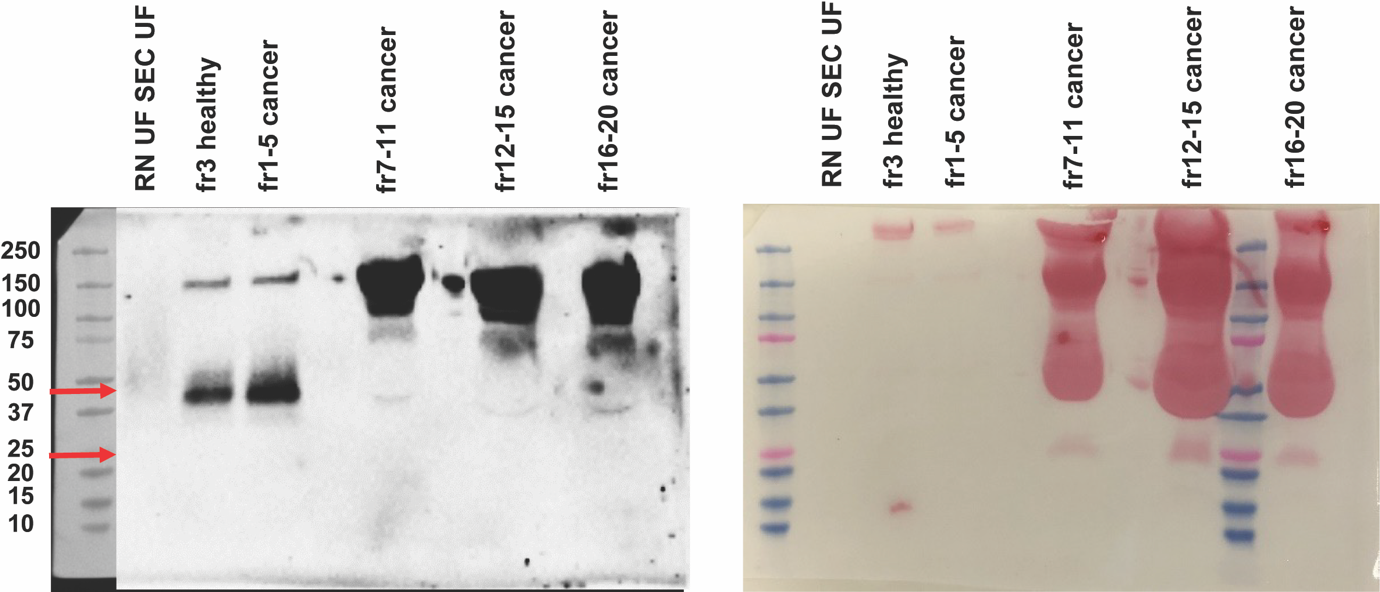

**Supplementary Figure 2:** Western blot with CD63 and CD81 labelling.

Left panel shows a western blot with samples from fractions of Code53 patient’s plasma. 40 ul UF-SEC-UF or AFC fraction was mixed with non-reducing sample buffer and run on a 4-15% gradient gel (Bio-Rad) for 45 minutes at 120V. Proteins were transferred onto a nitrocellulose membrane for 67 minutes at 250 mA. Right panel shows a Ponceau staining which was performed to check the transfer of proteins onto the membrane and shows a large amount of protein in later AFC fractions as expected. Membrane was first probed for mouse anti-CD63 (BD; 1:300, upper red arrow), followed by mouse-antiCD81 (BD; 1:500, bottom red arrow for expected height). No CD81 detected (1:500) after 10 minutes exposure.

**Supplementary Figure 3:**

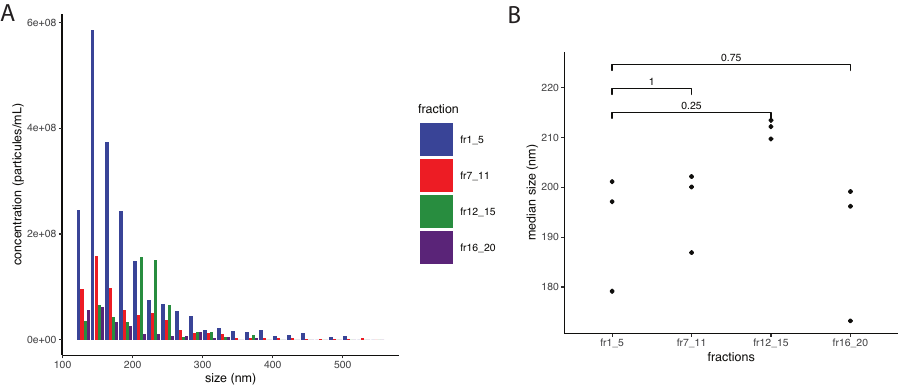

**Supplementary Figure 3:** concentration and sizes of particles in the different AFC fractions from one lung cancer patient (Code53) measured by Exoid (Izon). A: detailed particules concentration depending on their size. B: median size of particules depending on the fractions.

**Supplementary Figure 4:**

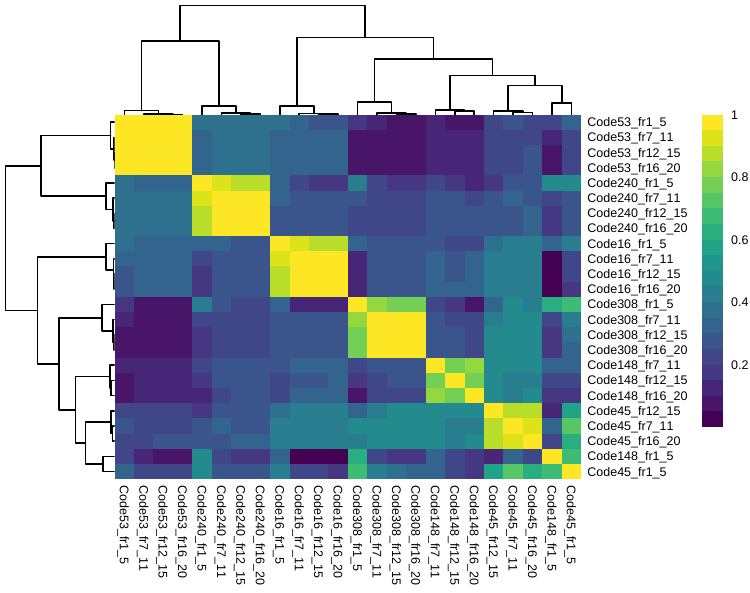

**Supplementary Figure 4: heatmap of the cross-correlation between the log_2_ratio copy number aberration profiles obtained from the lcWGS data.** The color scale indicates the correlation score (yellow=high, blue=low, Pearson correlation). Unsupervised clustering is applied to identify samples showing high correlation in their copy number aberration profiles.

**Supplementary Figure 5:**

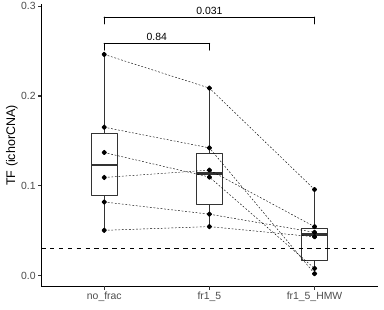

**Supplementary Figure 5: tumor fraction calculated from the SCNA data in cfDNA, short and long EV DNA.** Tumor fractions are determined using ichorCNA. Fr1_5_HMW: long EV DNA (>1000 bp) recovered from the AFC fraction 1-5. P values are indicated (paired Wilcoxon test).

**Supplementary Figure 6:**

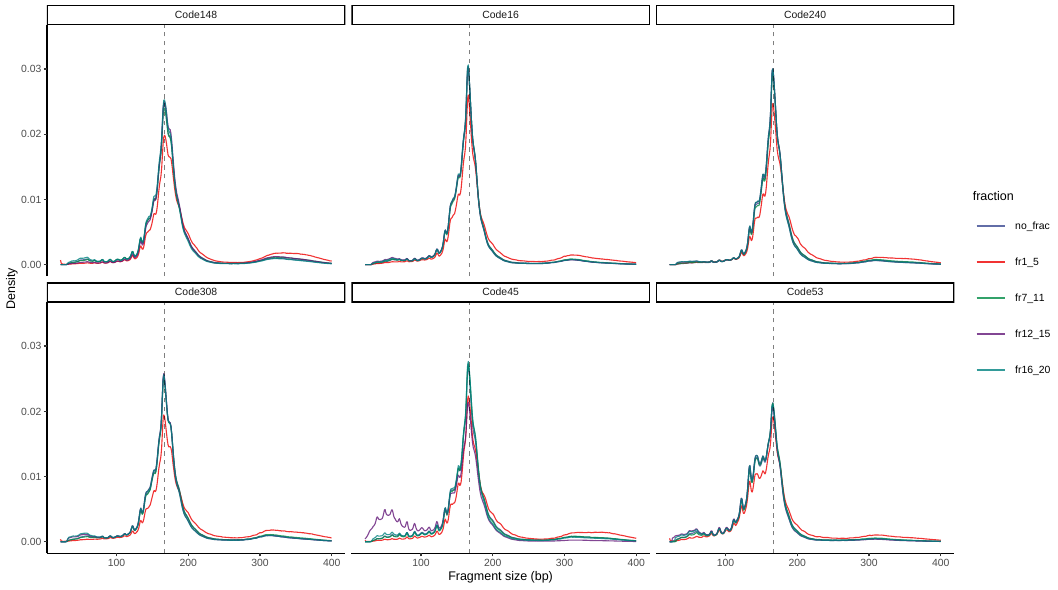

**Supplementary Figure 6: fragment size distribution of the cfDNA unfractionated samples (no_frac) and AFC samples (fr1_5; fr7_11; fr12_15; fr16_20).** Fragment sizes are recovered using paired-end lcWGS data.

**Supplementary Figure 7:**

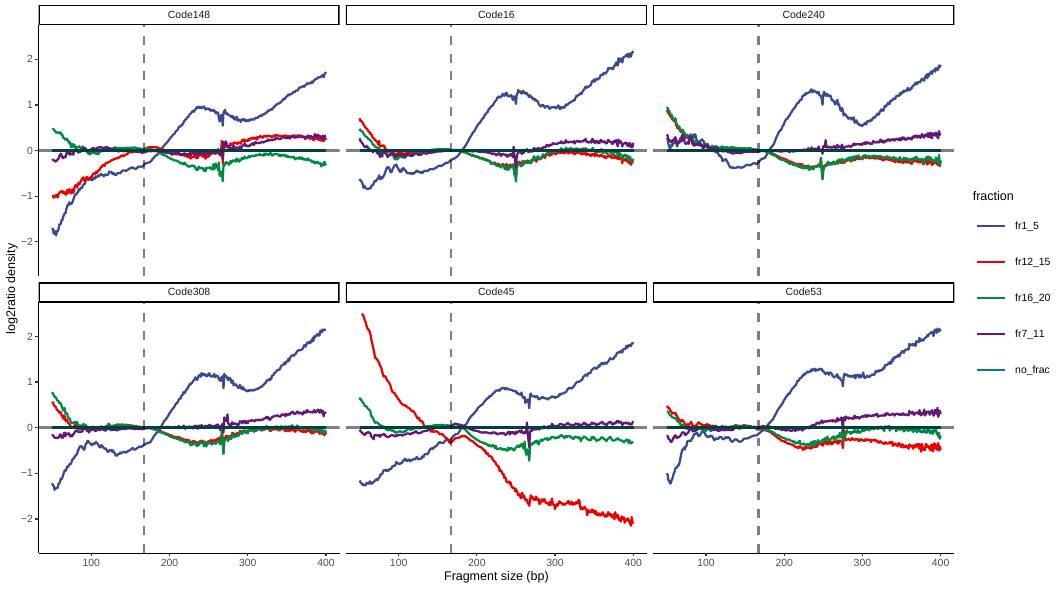

**Supplementary Figure 7: log_2_ratio of the fragment size distribution comparing the AFC samples (fr1_5; fr7_11; fr12_15; fr16_20) to the cfDNA size profile (no_frac).** Fragment sizes are recovered using paired-end lcWGS data.
